# The Impact of Education on Myopia: A bidirectional Mendelian randomisation analysis in UK Biobank

**DOI:** 10.1101/172247

**Authors:** Edward J. Mountjoy, Neil M. Davies, Denis Plotnikov, George Davey Smith, Santiago Rodriguez, Cathy Williams, Jeremy A. Guggenheim, Denize Atan

## Abstract

Myopia, or short-sightedness, is one of the leading causes of visual disability in the World. The prevalence of myopia has risen steadily over recent decades, reaching epidemic levels in Southeast Asia. Observational studies have reported associations between educational attainment and myopia. Whether education causes myopia, myopic children are more intelligent, or another factor, like higher socioeconomic status, causes both is unclear since observational studies are prone to confounding and randomised trials of education are unethical. Using bidirectional Mendelian Randomisation, a form of instrumental variable (IV) analysis free from confounding, we show that every additional year in education leads to an increase in myopic refractive error, but that myopia does not lead to higher educational attainment. Our results suggest that current educational methods contribute to the global burden of myopia, and argue that educational policies and practices should take account of this to reduce future visual disability in the population.

## Main text

Myopia, or short-sightedness, is one of leading causes of visual disability globally and the prevalence is rising rapidly^1,2^. Presently, ~50% of adults in the United States and Europe are myopic, with epidemic levels reported in Singapore, South Korea, China and other Asian communities^1-4^. Based on existing trends, the number of individuals affected by myopia worldwide is expected to rise from 1.4 billion currently, to 5 billion by 2050^4^.

Myopia is a refractive defect of the eye causing blurred distance vision as light is focussed in front of the retina instead of directly onto it; either because the refractive power of the cornea and lens is too great or the axial length of the eye is too long. Myopia can be corrected with spectacles, contact lenses or refractive surgery, but the risk of complications from potentially blinding conditions like retinal detachment, glaucoma and myopic maculopathy, increases with the longer axial lengths associated with high myopia^5-7^.

Myopia is highly heritable with >50 genetic variants identified by genome-wide association studies (GWAS)^8-10^. However, early visual experiences can affect ocular growth^11^, and the rapid rise in myopia prevalence over 1-2 generations implies important roles for environmental factors in causing myopia. Measures of education have long been linked with myopia^12,13^ and, conversely, the protective effect of time spent outdoors on myopia has been reported^14^. Yet, causal evidence for a role of education in myopia is lacking^3,15-18^ since randomised trials of education in children would be unethical.

Mendelian randomisation (MR) is a form of instrumental variable analysis^19^ that uses genetic variants as proxies for environmental exposures. It is an approach designed to reduce bias from confounding and reverse causation, to which observational epidemiology studies are susceptible (Figure 1). Given that genotypes are randomly assigned at conception, MR has been likened to a randomised trial by genotype^20,21^. The recent availability of large-scale GWAS data for education^22^ and myopia^10^ together with the genotypes of 488,377 individuals in the UK Biobank provided an opportunity to investigate the relationship between education and myopia by bidirectional MR analyses^23^ with high statistical power.

**Figure 1.**
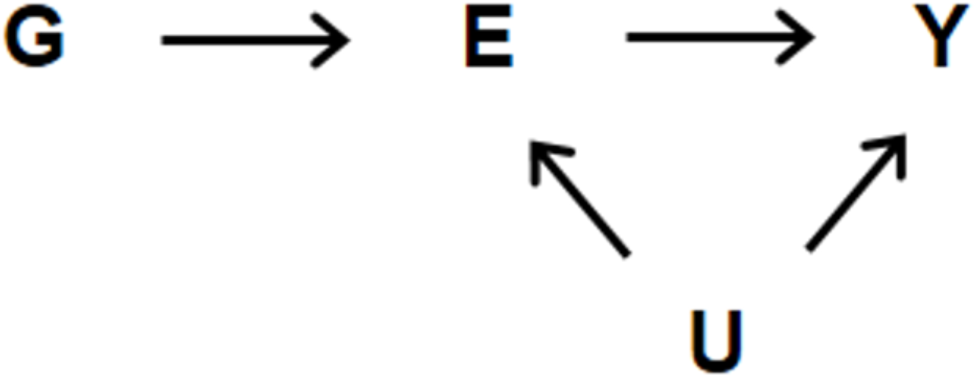
Principles of Mendelian Randomisation. Observational studies of the relationship of interest between an exposure *E* and an outcome *Y* may suffer bias due to confounder variables *U*. In Mendelian Randomisation, the effects of one or more specific genetic variants *G* on variables *E* and *Y* (i.e. *G* → *E* and *G* → *Y*) are measured in order to obtain an estimate of *E* → *Y* that is free from confounding by *U*, subject to certain assumptions.

In total, 69,798 UK Biobank participants had measures of educational attainment, refractive error and genome wide genetic data that passed validation (Extended Data Figure 1). In agreement with previous studies^3,17,18^, we found through standard ordinary least squares (OLS) observational analyses that more highly educated individuals were more myopic, i.e. they had more negative refractive errors (Table 1 & Extended Data Table 1). This relationship was linear at approximately −0.25D/year for those leaving full-time education from age 15-18 years; then reduced to approximately −0.10D/year for those leaving full-time education after age 18 (Figure 2). On average, every additional year spent in education was associated with a more myopic refractive error of −0.18D/year (95%CI: −0.19 to −0.17, p<2e-16). This association was largely unaffected by adjustment for measured confounders, including socioeconomic status (*Townsend Deprivation Index*, *birth weight*, and *breastfeeding during infancy*) and place of birth (*northing* and *easting* coordinates) (Table 1).

**Figure 2.**
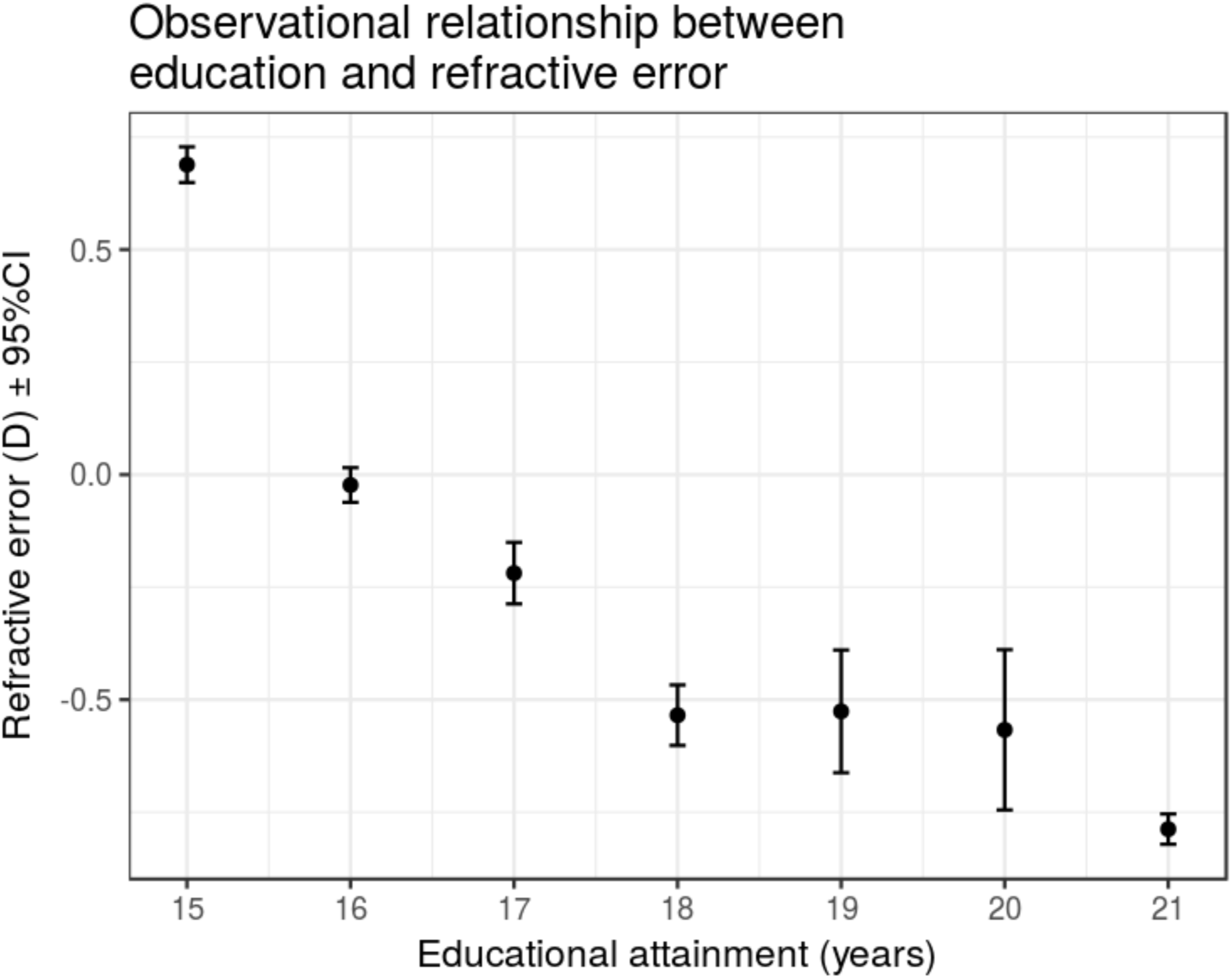
Observational association between educational attainment and refractive error. Graph of refractive error (y-axis) against age completed full-time education (x-axis) for 69,798 individuals in UK Biobank. On average, more educated individuals had higher levels of myopia.

**Table 1.**
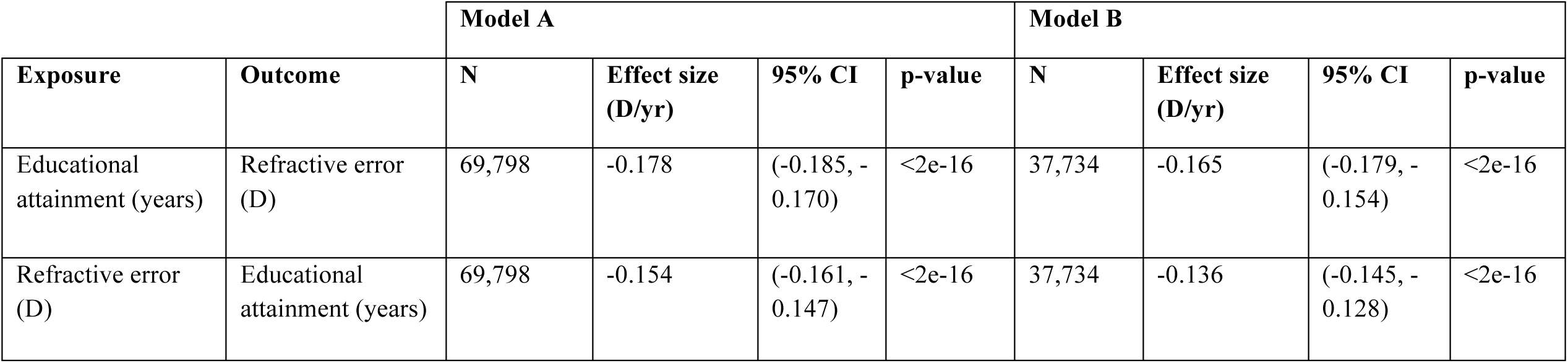
Observational association between educational attainment and refractive error. *Educational attainment* was defined as the age full time education was completed and *refractive error* was defined as the average measured mean spherical equivalent refractive error (aveMSE). Model A included *sex* and *age* as covariates. Model B included *age, sex, Townsend Deprivation Index (TDI), birth weight, whether breastfed, northing* and *easting* coordinates.

| Exposure | Outcome | Model A |  |  |  | Model B |  |  |  |
| --- | --- | --- | --- | --- | --- | --- | --- | --- | --- |
|  |  | N | Effect size (D/yr) | 95% CI | p-value | N | Effect size (D/yr) | 95% CI | p-value |
| Educational attainment (years) | Refractive error (D) | 69,798 | -0.178 | (-0.185, -0.170) | <2e-16 | 37,734 | -0.165 | (-0.179, -0.154) | <2e-16 |
| Refractive error (D) | Educational attainment (years) | 69,798 | -0.154 | (-0.161, -0.147) | <2e-16 | 37,734 | -0.136 | (-0.145, -0.128) | <2e-16 |

We next used bidirectional MR to assess the causality and direction of the association between educational attainment and refractive error. Bidirectional MR analyses consist of two separate MR calculations - one in each direction. Firstly, the causal effect of myopia on educational attainment was calculated using a weighted myopia allele score as the IV. Secondly, the causal effect in the opposite direction was calculated using a weighted education allele score as the IV. The allele score for myopia was derived from the most strongly associated genetic variants reported by Pickrell *et al*^10^ in a GWAS of self-reported myopia (N=191,843). Likewise, the allele score for educational attainment was derived from the genetic variants identified by Okbay *et al*^22^ in a large meta-analysis of GWAS of individuals of European descent (N=293,723) (Extended Data Table 3). These genetic variants were selected to use as our IVs because of their robust association with education and myopia, allowing us to construct strong instruments for making MR inferences. Thus, using the allele score for educational attainment as the IV, MR showed that every additional year spent in education resulted in a more myopic refractive error of −0.27 D/year (95%CI, −0.37 to −0.17, p=4.2e-8) (Table 2; Figure 3). The MR effect estimate was even greater in magnitude than the observational estimate (−0.27 vs. −0.18 D) suggesting that unmeasured confounders may have attenuated the latter relationship. Conversely, using the myopia allele score as the IV in MR provided no evidence that refractive error affected educational attainment (β_IV_ = −0.008 years/D, 95% CI −0.041 to 0.025, p=0.625) (Table 2; Figure 3). With our sample size of N=69,798, we had 80% power to detect an effect of education on refractive error ≥0.26D/yr. In the reciprocal direction, we had 80% power to detect an effect ≥0.048yr/D (Extended Data Figure 2), suggesting that our study had sufficient power to detect an effect of myopia on education, if present.

**Table 2.** Causal association between educational attainment and refractive error. Results of conventional multivariate linear regression and bidirectional MR. All regressions included *age* and *sex* as covariates. Abbreviation: DWH = Durbin-Wu-Hausman test for endogeneity.

| Exposure | Outcome | N | Observational estimate (OLS) |  | MR regression |  |  |  |  |  |
| --- | --- | --- | --- | --- | --- | --- | --- | --- | --- | --- |
|  |  |  | Effect size | 95% CI | F-stat | Partial R<sup>2</sup> | p-value (DWH) | Effect size | 95% CI | p-value |
| **Educational attainment** | **Refractive error** | 69,798 | -0.178 D/yr | (-0.185, -0.170) | 495.4 | 0.71% | 0.0589 | -0.270 D/yr | (-0.368, -0.173) | 4.2e-8 |
| **Refractive error** | **Educational attainment** | 69,798 | -0.154 yr/D | (-0.161, -0.147) | 3154.6 | 4.32% | 4.05e-19 | -0.008 yr/D | (-0.041, 0.025) | 0.625 |

**Figure 3.**
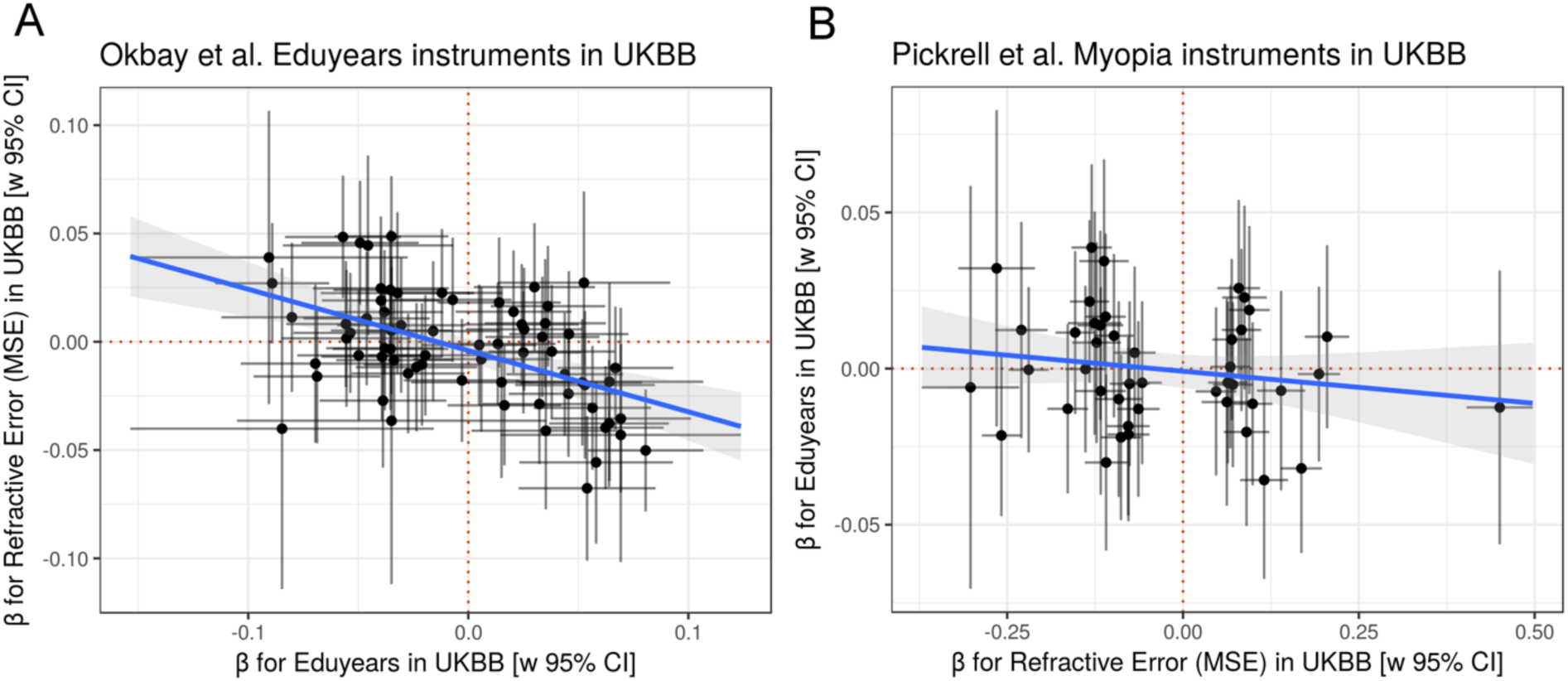
Association of educational attainment and myopia instrumental variables identified in independent cohorts with educational attainment and refractive error in UK Biobank. **(A)** 69 variants associated with educational attainment in Okbay *et al* (2016) were linked to higher levels of myopia in UK Biobank; **(B)** 44 variants associated with myopia in Pickrell *et al* (2016) did not lead to higher levels of educational attainment in UK Biobank. Regression line and standard errors fitted using robust linear regression.

MR analyses are based on two pertinent assumptions: (i) the genetic instruments are only associated with the outcome via the exposure, and (ii) the genetic instruments are not associated with any confounders of the exposure-outcome relationship. The Durbin-Wu-Hausman (DWH) test for endogeneity is a method to check for endogenous variables in a regression model that would create bias, e.g. from omitted variables, measurement error or reverse causation^25,26^. There was weak evidence that the IV estimate using the education allele score differed from the observational point estimate (DWH-p=0.059), with the IV estimate suggesting a larger negative association (Table 2). There was strong evidence that IV estimate using the myopia allele score was a departure from the observational point estimate (DWH-p=4.05e-19) (Table 2).

The second core assumption of MR is that the IV is not associated with a confounder of the exposure-outcome association^20^. There was evidence that the geographical co-ordinate *northing* (measured northward distance in UK) is a confounder of the education-refractive error relationship; *northing* was negatively associated with education (β = −1.61e^−6^, 95% CI, −1.78e^−6^ to −1.45e^−6^) and positively with refractive error (β = 1.16e^−6^, 95% CI 9.84e^−7^ to 1.33e^−6^). *Northing* was also associated with the education (p=6.8e-5) and myopia (p=6.1e-3) allele scores (Extended Data Table 2). Compared to standard regression, confounding bias plots suggest that inclusion of the *northing* variable in the IV analysis may result in a greater degree of bias for the education allele score (Figure 4A) but not for the myopia allele score (Figure 4B).

**Figure 4.**
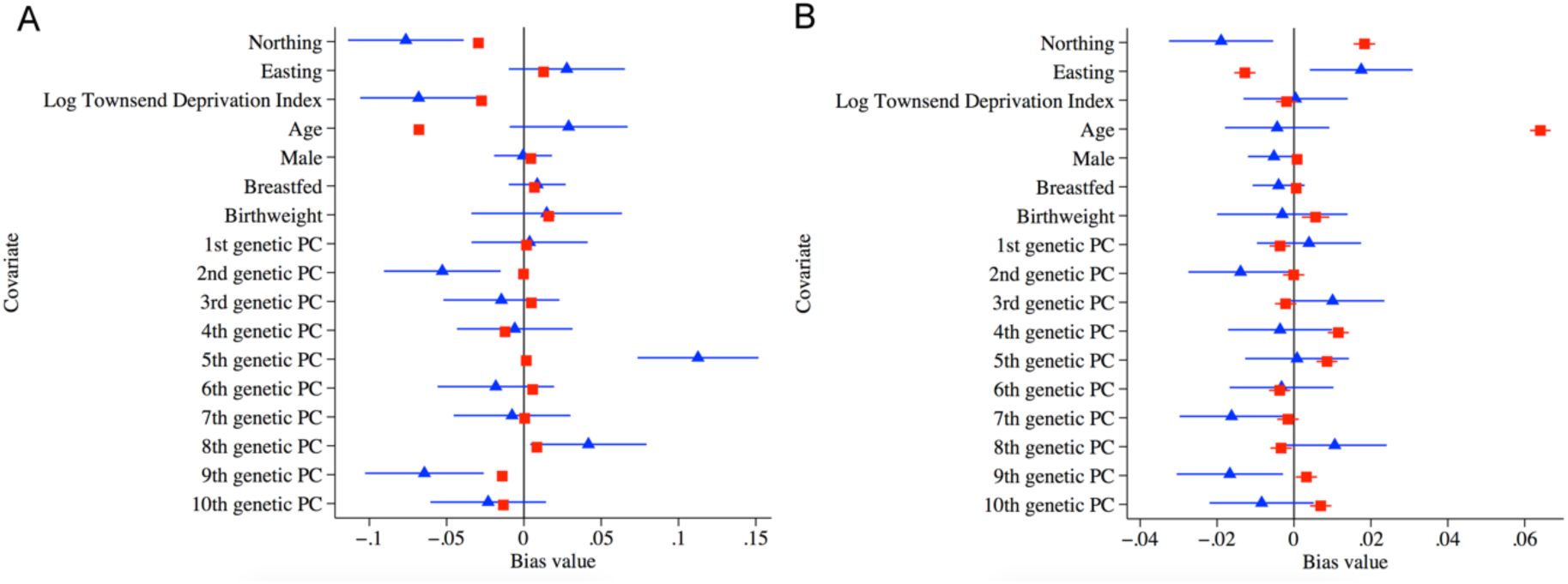
Confounding bias plots. Plots showing the relative bias in the instrumental variable estimate (blue) and standard multivariate regression estimate (red) when **(A)** estimating the effect of educational attainment on refractive error; and **(B)** estimating the effect of refractive error on educational attainment.

In contrast, the geographical *easting* coordinate was positively associated with education (β = 8.90e^−7^, 95% CI, 6.82e^−7^ to 1.10e^−6^) and negatively associated with refractive error (β = −1.03e^−6^, 95% CI, −1.25e^−6^ to −8.06e^−6^). It was weakly associated with the myopia allele score (p=0.01). However, there no evidence to suggest a greater degree of bias in the IV analysis compared to a standard regression with the inclusion of the *easting* variable (Figure 4B). We identified one further confounding variable as population stratification principle component 9, which incurred a greater degree of bias in the IV regression compared to OLS regression. While these confounders (*northing* and population stratification component 9) may bias our MR effect estimates, we restricted our analyses to unrelated individuals of European ancestry to limit the likelihood of major bias from population stratification.

MR causal estimates were also calculated using the MR-Egger and Weighted Median methods (which partially relax the assumptions required for MR causal estimates to be valid) as well as alternative methods of integrating IV estimates across individual SNPs. All of these approaches yielded causal estimates indicating that increasing levels of education led to a more myopic refractive error (by −0.22 to −0.44 D/year; p<0.05 for all methods), while there was no evidence that a more myopic refractive error led to greater educational attainment (p>0.05 for all methods) (Extended Data Table 4). An advantage of MR-Egger is that it gives a valid test for causality even if invalid instruments are used, e.g. due to horizontal pleiotropy^27^. The ubiquity of exposure to education in populations with available genotype data means that it is not possible to assess individuals who are completely free of our outcome, specifically education. With MR-Egger, a deviation of the intercept estimate from zero would suggest the existence of horizontal pleiotropy, i.e. where certain genetic variants affect the outcome via a different biological pathway from the exposure under investigation. In practice, there was no evidence that the Egger intercept deviated from zero either for education causing refractive error (intercept=0.007, SE=0.006, p=0.22) or refractive error causing education (intercept=−0.002, SE=0.007, p=0.81), indicating that there was no evidence for horizontal pleiotropy. However, such bias cannot be ruled out definitively until we gain more knowledge of the mechanisms by which these genetic variants affect the traits described here.

To ensure that the association between educational attainment and myopia was not an artefact of the non-normal distribution of the variable, *age completed schooling*, educational attainment was recoded using two alternative methods: (1) dichotomising the cohort into participants who finished their schooling when >16 years versus £16 years of age; and (2) excluding individuals who had attended college or university. Encoding education as a dichotomous trait (>16 years vs <16 years of age when completed schooling) produced the same pattern of causality as the continuous variable, *age completed schooling*; i.e. education had an effect on refractive error (β_IV_ = −0.347 D/LOD(education), 95% CI −0.482 to −0.220) while refractive error did not have an effect on education (β_IV_ = −0.0004 LOD(education)/D, 95% CI −0.028 to 0.028) (Extended Data Table 5).

When individuals who had attended university or college were excluded from the analyses, there was a similar point estimate of the effect of education on refractive error (β_IV_ = −0.228 D/yr, 95% CI −0.479 to 0.018, p=0.066) with larger standard errors. This is attributable, in part, to the reduced sample size (N=45,535). Again, there was no evidence that refractive error had an effect on educational attainment (β_IV_ = −0.004 yr/D, 95% CI −0.035 to 0.027, p=0.80) (Extended Data Table 5).

MR is a powerful approach for testing causal hypotheses in epidemiology^28,29^. The large sample size and robustly-associated genetic instruments used here meant that causal effects could be estimated with high precision. Our findings are in agreement with the single previous study to address the causal relationship between education and myopia, a meta-analysis of 3 European-ancestry cohorts (N=5,649) using MR^30^. Cuellar-Partida *et al.*^30^ included 17,749 SNPs to construct their polygenic risk score as an IV for educational attainment. The authors reported that each year of educational attainment led to a more myopic refractive error of −0.46 D/year (p=1.0e-3). However, the study by Cuellar-Partida *et al*.^30^ was underpowered, which may explain why results from 2 of the 3 cohorts they studied were non-significant. Furthermore, their methodology risked violating one of the key assumptions of MR; firstly because the thousands of SNPs used to create their IV may well have included pleiotropic variants with direct effects on both educational attainment and refractive error; and secondly, because there were likely to be SNPs associated with educational attainment that were in LD with refractive error variants. The much larger sample size in our study permitted the use of less than 100 variants to use as IVs for educational attainment and refractive error. Thus, we were able to mitigate against the risk of LD between the major risk variants for the two traits explaining the underlying associations between education and myopia. Crucially, our analyses provided strong evidence that this relationship arose from a *causal* effect of education on refractive error, and not via reverse causation or confounding by influences such as socioeconomic status.

Myopes, by definition, have better near vision than distance vision and require less accommodative effort for near work and study, and so myopia has been proposed as an educational advantage^31^. Despite the general perception that myopes are more studious than non-myopes, we found no evidence that refractive error affected educational attainment.

Conversely, known environmental risk factors for myopia provide intriguing clues about how educational exposure may influence refractive error. Randomised controlled trials have demonstrated that time spent outdoors during childhood protects against myopia development, at least in part^14,32^; whereas the time that children spend doing near work activities, such as reading, has been less consistently linked to the incidence and progression of myopia^15,33-35^. Indeed, the time that children spend outdoors is typically independent of their near work activities, as measures of the two are generally uncorrelated^36-38^. Myopic children tend to engage in less physical activity, such as sports^38,39^. Individuals who spend more years in education may spend a larger proportion of their time indoors, so it is plausible that spending less time outdoors may be one mechanism through which higher levels of educational attainment cause myopia; but this requires further investigation.

Given the rapid rise in the global prevalence of myopia and the economic burden of myopia and its vision-threatening complications, our findings have important implications for educational practices. Educational policy makers should be aware that the current practices employed for educating our children to promote personal and economic health may have the unintended consequence of causing increasing levels of myopia and later visual disability as a result.

## Methods

### The study cohort in UK Biobank

We analysed cross-sectional data from the baseline assessment of the UK Biobank project^40^. During the period 2006 to 2010, UK Biobank recruited 502,649 participants aged 37 to 73 years-old. Participants attended 1 of 22 assessment centres across the UK, at which they completed a touch-key questionnaire, had a face-to-face interview with a trained nurse, and underwent physical assessments. All participants completed sociodemographic questionnaires, which included questions on past educational or professional qualifications. Towards the end of the UK Biobank recruitment exercise, a detailed ophthalmic assessment was introduced. Approximately 23% of participants underwent the ophthalmic assessment. Participants who had withdrawn consent were excluded. A total of 69,798 participants had valid education, refractive error and genetic data available (Extended Data Figure 1).

### (i) The definition of educational attainment in UK Biobank

Educational attainment was defined by the question *age completed full-time education* (variable 845) in UK Biobank. The question was ascertained only for participants that did not have a college or university degree (variable 6138, answer 1). Following Okbay *et al.*, participants with a college or university degree were coded as having left full-time education at the age of 21. Participants who reported their *age completed full-time education* was less than 15 years were assigned a value of 15 years. As schooling systems differ between countries, only participants born in England, Scotland or Wales were included in the analyses (variables 1647, 20115).

### (ii) The definition of refractive error in UK Biobank

Refractive error was measured as part of the ophthalmic assessment using non-cycloplegic autorefraction (Tomey RC5000 autorefractor; performed after removal of habitual spectacles or contact lenses). Up to 10 repeat measurements were taken. Measurements were excluded if the autorefractor reading was flagged as unreliable (variables 5090/5091). Spherical power (variables 5085/5084) and cylindrical power (variables 5086/5087) were averaged over repeat measurements. Mean spherical equivalent (MSE) refractive error for each eye was calculated (*Spherical power* + 0.5 * *Cylindrical power*). The mean of the left and right MSE (aveMSE) was taken as the participant’s refractive error and used in our analyses (N=127,412). For participants with repeat measurements from separate visits (baseline visit and subsequent visits), only the baseline measurement was used. Individuals with preexisting eye conditions that could affect refractive error were excluded from the analyses, namely: cataracts (variables 6148, 5324, 5441), refractive laser eye surgery (variable 5325), injury or trauma resulting in vision loss (variable 5419), or corneal graft surgery (variable 5328). A total of 10,984 individuals with pre-existing eye conditions were excluded.

### (iii) The genotype data in UK Biobank

Participants were genotyped using one of two platforms: the Affymetrix UK BiLEVE Axiom array or the Affymetrix UK Biobank Axiom array. The genetic data underwent rigorous quality control procedures and was phased and imputed against a reference panel of Haplotype Reference Consortium (HRC), UK10K and 1000 Genomes Phase 3 haplotypes^41^. Due to an issue with the imputation of UK10K and 1000 Genomes variants, analysis has been restricted to HRC variants only. Samples were excluded based on the following genotype-based criteria: non-European ancestry, relatedness, mismatch between genetic sex and self-reported gender, putative aneuploidy (variable 22019), outlying heterozygosity, and excessive missingness (variable 22027)^41^.

## Statistical Methods

### (i) Ordinary Least Squares (OLS) observational analyses

Observational associations between refractive error and educational attainment were assessed using linear regression adjusted for *sex* and *age*. The regression was then repeated with adjustment for additional potentially confounding variables: *Townsend deprivation index (TDI), birth weight, whether breastfed*, and geographic coordinates of place of birth rounded to the nearest kilometre (*northing* and *easting* coordinates).

### (ii) Mendelian Randomisation

#### (a) The genetic instruments used for Mendelian Randomisation

Pickrell *et al*^10^ reported the results of a GWAS for self-reported myopia in a sample of N=191,843 individuals (106,086 cases, 85,757 controls) carried out by the personal genomics company 23andMe. The 50 most strongly associated variants were reported. Six variants (rs5022942, rs10887265, rs71041628, rs34016308, rs11658305 and rs201140091) were not in the HRC panel, leaving 44 for use as genetic instruments in our analysis.

Okbay *et al*^22^ identified 74 variants associated with educational attainment in a large meta-analysis of GWAS of individuals of European descent (N=293,723). Educational attainment was defined as the number of years spent in schooling. Okbay *et al* used UK Biobank as a replication cohort. Therefore, we only used genetic variants and summary statistics from the discovery analysis by Okbay *et al* (available at: http://ssgac.org/documents/EduYears_Discovery_5000.txt [accessed 30/03/2017]). Five variants (rs9320913, rs148734725, rs62056842, rs114598875, rs8005528) were not in the HRC panel, leaving 69 variants for use as instruments.

These genetic variants are described in Extended Data Table 3.

#### (b) The generation of allele scores for Mendelian Randomisation

Multiple genetic variants were combined into a single weighted allele score for each trait. An allele score, compared to individual variants, has been shown to improve the coverage properties and reduce the bias of instrumental variable (IV) estimates^42^. Effect size estimates from the original GWAS publications were used to weight variants when constructing allele scores. Variants were harmonised with UK Biobank to ensure correct coding of the effect allele. Genotype probabilities were converted to effect allele (*a*) and non-effect allele (*A*) dosages. Allele scores were calculated by summing the product of the weights and dosages across all *n* variants:

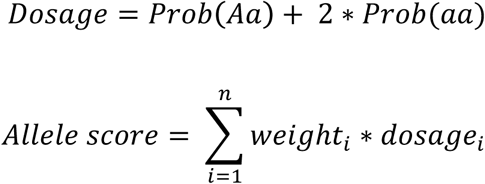

The proportion of variance in the phenotype variable explained by the allele score IV was calculated by regressing the phenotype on its respective allele score. The myopia allele score explained 4.32% (F=3153) of the variance in aveMSE. The education allele score explained 0.71 % (F=495) of the variance in educational attainment in UK Biobank. The large F-statistics suggest the analysis should not be affected by weak instrument bias.

#### (c) The implementation of MR

MR was implemented using the two-stage least squares method in the R package *ivpack*^43^. *Age* and *sex* were included as covariates. The strength of association in the first stage regressions between allele score and exposure were assessed with F-tests, to assess the risk of weak instrument bias^44^. Effect estimates from the observational analysis and second-stage instrumental analysis were tested for endogeneity using the Durbin-Wu-Hausman (DWH) test^25,26^. Statistical power was assessed using the mRnd online calculator^45^ (available: http://cnsgenomics.com/shiny/mRnd/).

#### (d) Sensitivity analyses for confounding, pleiotropy and artefact

MR analyses assume that covariates are randomly distributed with respect to genotype^20^. A number of variables were found to confound the education-refractive error relationship and were associated with one or both instrumental variables (allele scores) (Extended Data Table 2). Confounding bias plots^46,47^ were used to assess relative bias in the IV estimate compared to standard multivariate regression. Additionally, suspected confounding factors were included as covariates in supplementary analyses (Extended Data Table 5).

Under a second assumption of MR, genetic variants with pleiotropic effects are invalid instruments. This can be problematic when genetic variants are used without regard for the biological mechanisms through which they affect the exposure. MR-Egger is a method that gives a valid test for causality despite the presence of invalid instruments^27^. With MR-Egger regression, deviation of the intercept estimate from zero suggests the existence of horizontal pleiotropy. However, MR-Egger is not valid for one-sample MR, used for the main analyses in this study, as the error terms for the genetic variant (G)-exposure (E) and G-outcome (O) associations will be correlated. It was, therefore, necessary to run MR-Egger as a split sample analysis, whereby the sample was randomly split in half, then G-E and G-O associations were calculated separately in the independent groups (Extended Data Table 4, sheets 2 & 3). MR-Egger was implemented using the R package MRBase^48^ (available: github.com/MRCIEU/TwoSampleMR).

Finally, to ensure the association between educational attainment and myopia was not an artefact of the non-normal distribution of the variable *age completed schooling*, educational attainment was recoded using two alternative methods: (1) dichotomisation into age >16 years when completed schooling and age £16 years when completed schooling; and (2) excluding individuals who attended college or university. The results were compared with the original analyses using the continuous variable *age completed schooling*.

## Code availability

Code used to run the analysis is available on GitHub: https://github.com/edm1/myopia-education-MR

## Acknowledgements

This research has been conducted using the UK Biobank Resource (application #8786). The research was supported by funding from: the Medical Research Council (MRC), Bristol Centre for Systems Biomedicine (SD 1221), the Global Education Program of the Russian Federation government (DP), National Eye Research Centre grant SAC015 (JAG, CW), an NIHR Senior Research Fellowship award SRF-2015-08-005 (CW), the MRC and the University of Bristol fund the MRC Integrative Epidemiology Unit [MC_UU_12013/1, MC_UU_12013/8, MC_UU_12013/9]. NMD is supported by the Economics and Social Research Council (ESRC) via a Future Research Leaders grant [ES/N000757/1].

## Author contributions

E.M. and N.D. analysed the data with assistance from G.D.S.; D.A., J.G., D. P., C.W. and S.R supervised the project. D.A. and J.G conceived the project and wrote the manuscript with E.M.

## Author information

The authors declare no competing financial interests. Correspondence and requests for materials should be addressed to J.G. or D.A.

## Data availability

The data used in this study can be accessed by contacting UK Biobank (www.ukbiobank.ac.uk). This analysis was approved by the UK Biobank access committee as part of project 8786. Consent was sought by UK Biobank as part of the recruitment process.

## Extended Data Figures and Tables

**Extended Data Figure 1.**
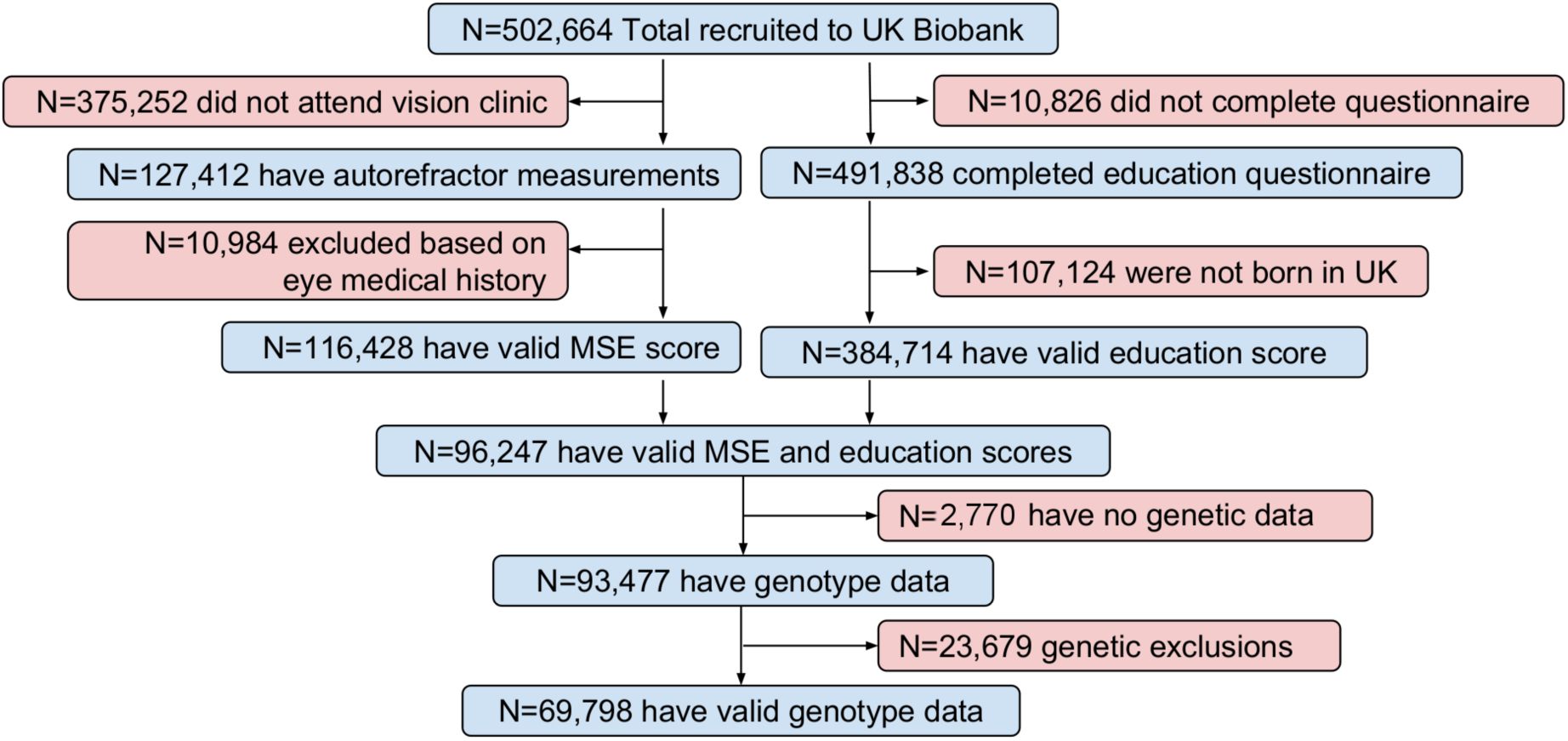
Numbers of participants in UK Biobank that passed validation for the MR study.

**Extended Data Figure 2.**
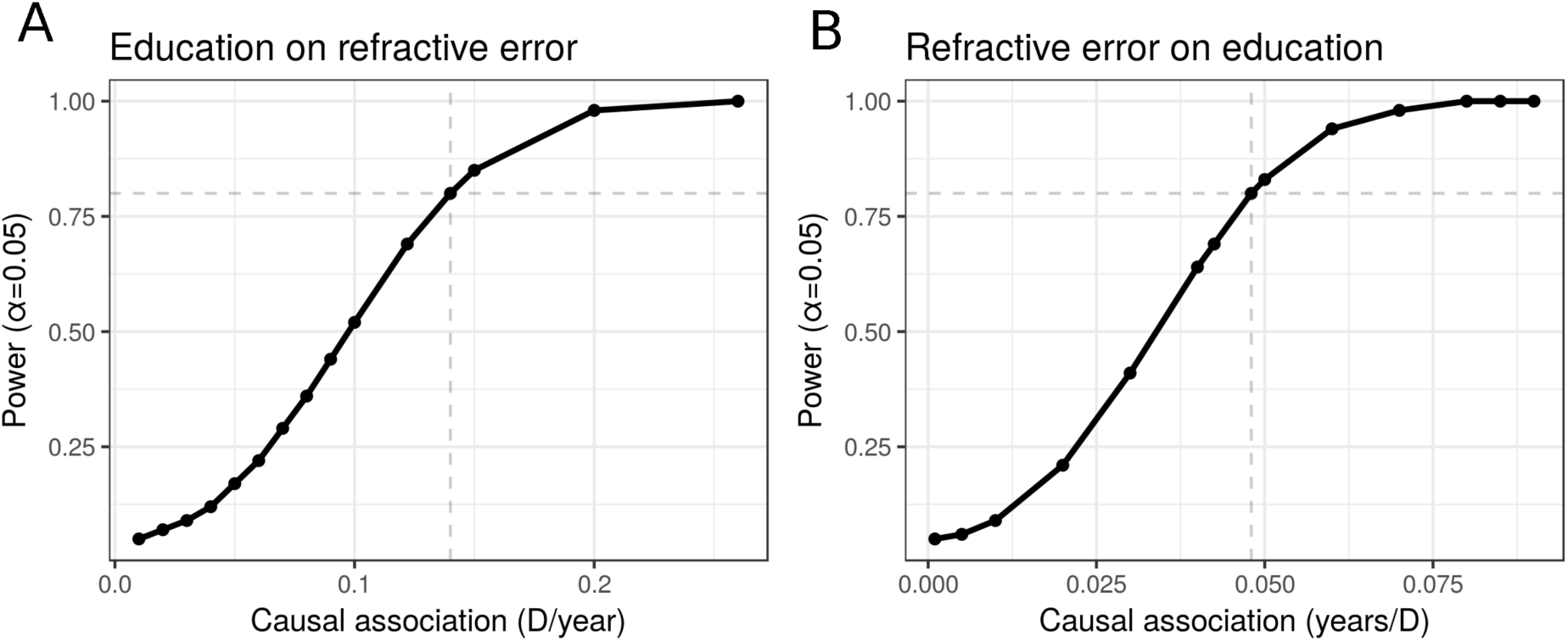
Mendelian Randomisation power calculation for the available sample size in UK Biobank (N=69,798). Causal association of: **(A)** education on refractive error using allele score with R^2^=0.0073 (80% power = 0.14 D/yr); and **(B)** refractive error on education using allele score with R^2^=0.0442 (80% power = 0.048 yr/D).

**Extended Data Table 1.**
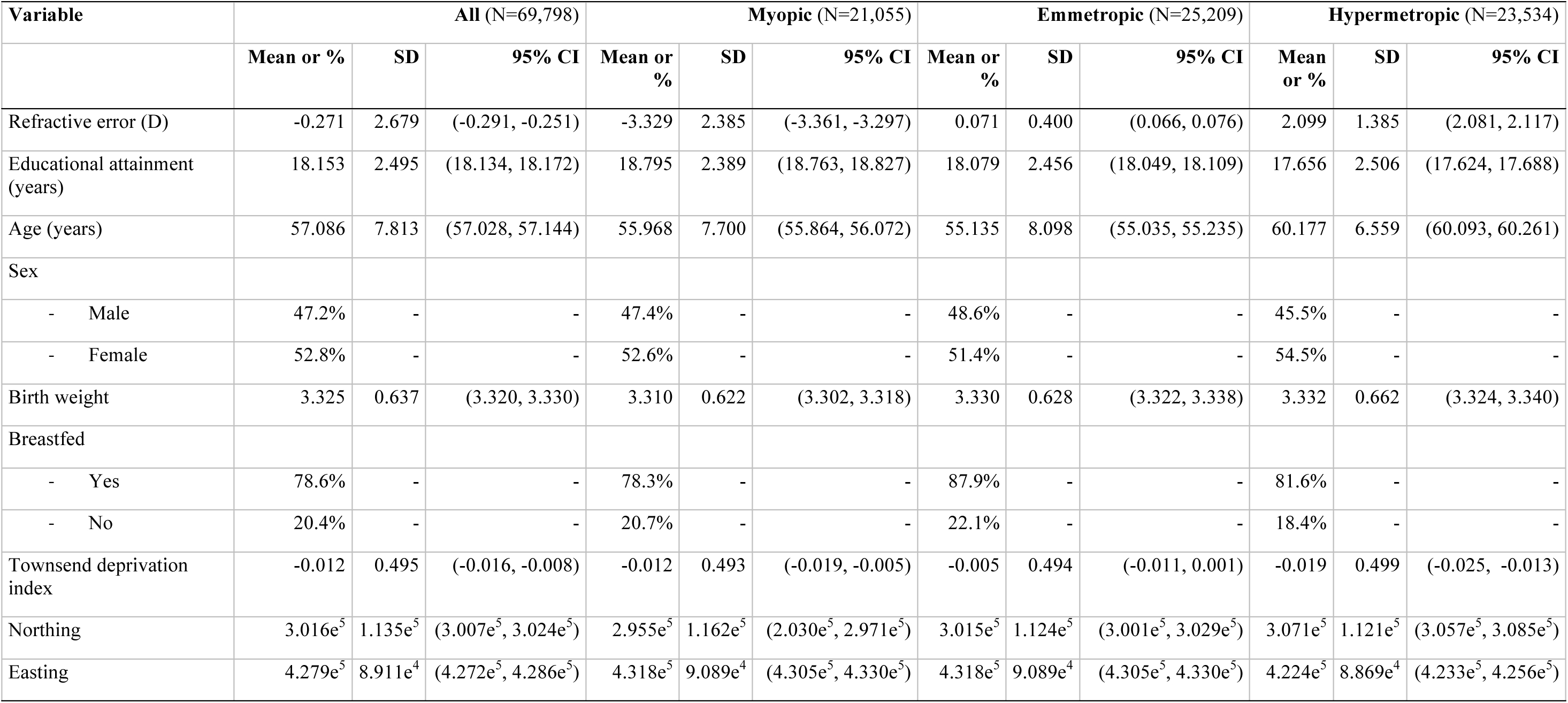
Baseline characteristics of UK Biobank cohort for refractive error, educational attainment and confounding variables. Refractive errors were defined as: (i) myopia ≤ 0.75; (ii) −0.75 < emmetropia < 0.75; (iii) hypermetropia > 0.75. *Townsend deprivation index* (TDI) was natural log-transformed.

| Variable | All (N=69,798) |  |  | Myopic (N=21,055) |  |  | Emmetropic (N=25,209) |  |  | Hypermetropic (N=23,534) |  |  |
| --- | --- | --- | --- | --- | --- | --- | --- | --- | --- | --- | --- | --- |
|  | Mean or % | SD | 95% CI | Mean or % | SD | 95% CI | Mean or % | SD | 95% CI | Mean or % | SD | 95% CI |
| Refractive error (D) | -0.271 | 2.679 | (-0.291, -0.251) | -3.329 | 2.385 | (-3.361, -3.297) | 0.071 | 0.400 | (0.066, 0.076) | 2.099 | 1.385 | (2.081, 2.117) |
| Educational attainment (years) | 18.153 | 2.495 | (18.134, 18.172) | 18.795 | 2.389 | (18.763, 18.827) | 18.079 | 2.456 | (18.049, 18.109) | 17.656 | 2.506 | (17.624, 17.688) |
| Age (years) | 57.086 | 7.813 | (57.028, 57.144) | 55.968 | 7.700 | (55.864, 56.072) | 55.135 | 8.098 | (55.035, 55.235) | 60.177 | 6.559 | (60.093, 60.261) |
| Sex |  |  |  |  |  |  |  |  |  |  |  |  |
| - Male | 47.2% | - | - | 47.4% | - | - | 48.6% | - | - | 45.5% | - | - |
| - Female | 52.8% | - | - | 52.6% | - | - | 51.4% | - | - | 54.5% | - | - |
| Birth weight | 3.325 | 0.637 | (3.320, 3.330) | 3.310 | 0.622 | (3.302, 3.318) | 3.330 | 0.628 | (3.322, 3.338) | 3.332 | 0.662 | (3.324, 3.340) |
| Breastfed |  |  |  |  |  |  |  |  |  |  |  |  |
| - Yes | 78.6% | - | - | 78.3% | - | - | 87.9% | - | - | 81.6% | - | - |
| - No | 20.4% | - | - | 20.7% | - | - | 22.1% | - | - | 18.4% | - | - |
| Townsend deprivation index | -0.012 | 0.495 | (-0.016, -0.008) | -0.012 | 0.493 | (-0.019, -0.005) | -0.005 | 0.494 | (-0.011, 0.001) | -0.019 | 0.499 | (-0.025, -0.013) |
| Northing | 3.016e<sup>5</sup> | 1.135e<sup>5</sup> | (3.007e<sup>5</sup>, 3.024e<sup>5</sup>) | 2.955e<sup>5</sup> | 1.162e<sup>5</sup> | (2.030e<sup>5</sup>, 2.971e<sup>5</sup>) | 3.015e<sup>5</sup> | 1.124e<sup>5</sup> | (3.001e<sup>5</sup>, 3.029e<sup>5</sup>) | 3.071e<sup>5</sup> | 1.121e<sup>5</sup> | (3.057e<sup>5</sup>, 3.085e<sup>5</sup>) |
| Easting | 4.279e<sup>5</sup> | 8.911e<sup>4</sup> | (4.272e<sup>5</sup>, 4.286e<sup>5</sup>) | 4.318e<sup>5</sup> | 9.089e<sup>4</sup> | (4.305e<sup>5</sup>, 4.330e<sup>5</sup>) | 4.318e<sup>5</sup> | 9.089e<sup>4</sup> | (4.305e<sup>5</sup>, 4.330e<sup>5</sup>) | 4.224e<sup>5</sup> | 8.869e<sup>4</sup> | (4.233e<sup>5</sup>, 4.256e<sup>5</sup>) |

**Extended Data Table 2. Observational (OLS) associations of confounding variables with education and myopia allele scores and outcomes.**

Variables that confound the education – refractive error relationship which are also associated with an allele score are highlighted in green.

(Extended_data_table_2.xlsx)

**Extended Data Table 3. Genetic variants from Okbay *et al* and Pickrell *et al* used to construct the education and myopia allele scores in this study.**

Association scores in the original study are shown in green; association scores in UK Biobank are shown in blue.

(Extended_data_table_3.xlsx)

**Extended Data Table 4. Causal estimates of education on refractive error and refractive error on education using methods implemented in MRBase and a split sample in UK Biobank.**

Sheet 1 shows estimates using various methods from MRBase. Sheets 2 & 3 show per variant associations with education and refractive error in split samples A (blue) and B (green).

(Extended_data_table_4.xlsx)

**Extended Data Table 5. Table of MR results from sensitivity analyses using alternative encodings of educational attainment and additional covariates.**

(Extended_data_table_5.xlsx)

